# Porous membrane electrical cell-substrate impedance spectroscopy for versatile assessment of biological barriers in vitro

**DOI:** 10.1101/2023.06.26.546641

**Authors:** Oleg Chebotarev, Alisa Ugodnikov, Craig A. Simmons

## Abstract

Cell culture models of endothelial and epithelial barriers typically use porous membrane inserts (e.g., Transwell inserts) as a permeable substrate on which barrier cells are grown, often in co-culture with other cell types on the opposite side of the membrane. Current methods to characterize barrier function in porous membrane inserts can disrupt the barrier or provide bulk measurements that cannot isolate barrier cell resistance alone. Electrical cell-substrate impedance sensing (ECIS) addresses these limitations but its implementation on porous membrane inserts has been limited by costly manufacturing and low sensitivity. Here we present porous membrane ECIS (PM-ECIS), a cost-effective method to adapt ECIS technology to porous substrate-based *in vitro* models. We demonstrate high fidelity patterning of electrodes on porous membranes that can be incorporated into well plates of a variety of sizes with excellent cell biocompatibility with mono- and co-culture set ups. PM-ECIS provided sensitive, real-time measurement of isolated changes in endothelial cell barrier impedance with cell growth and barrier disruption. Barrier function characterized by PM-ECIS resistance correlated well with permeability coefficients obtained from molecular tracer permeability assays performed on the same cultures, validating the device. Integration of ECIS into conventional porous cell culture inserts provides a versatile, sensitive, and automated alternative to current methods to measure barrier function in vitro, including molecular tracer assays and transepithelial/endothelial electrical resistance (TEER).

## 1. Introduction

Biological barriers play a key role in maintaining homeostasis and protection against infection, with tissue-specific endothelial or epithelial cells working in tandem with support cells and matrices to control transport of substances into compartments of the body (Cacopardo et al., 2019; Yeste et al., 2018). Beyond maintaining concentration gradients and excluding pathogens, biological barrier function is also critical in the absorption and distribution of pharmaceutical agents within different tissues (Cacopardo et al., 2019; Yang et al., 2017).

The most common in vitro model of biological barriers is the porous cell culture insert (e.g., Transwell insert), in which endothelial or epithelial barrier cells are grown on a permeable membrane that can be placed in a traditional cell culture well plate to create a cell monolayer as a barrier with luminal and abluminal compartments. This is a simple, robust method for emulating the multicellular environment of biological barriers. The blood-brain barrier (BBB), for example, is frequently modeled in this setup, enabling investigation of the effects of conditioned medium and various arrangements of supporting cells (e.g., astrocytes, pericytes) in direct or indirect contact with endothelial cells grown on porous cell culture inserts (Boveri et al., 2005; Cecchelli et al., 1999; Dohgu et al., 2005; Gaillard et al., 2001; Hatherell et al., 2011; Nakagawa et al., 2009, 2007; Perrière et al., 2007; Rubin et al., 1991).

Evaluation of barrier function in porous cell culture inserts often involves measurement of the diffusion of a molecular tracer (e.g., FITC-dextran) to determine barrier permeability (Stolwijk et al., 2015) However, the dyes used in these assays may themselves interfere with molecular transport function and barrier integrity (Srinivasan et al., 2015), thus motivating the use of electrical sensing as a non-invasive alternative. The most common electrical sensing technique for biological barriers is transepithelial/endothelial electrical resistance (TEER), which measures conductance of ions across paracellular junctions (Maoz et al., 2017) through the application of an alternating current by electrodes placed on either side of the membrane. In the standard TEER setup, AC current at a frequency of 12.5 Hz – selected to avoid charging the electrode and/or cell layer (Srinivasan et al., 2015) – is applied across the cell layer to determine barrier resistance. Because TEER calculations include normalization to culture area, it is theoretically possible to compare values between various barrier types and across different culture setups.

While TEER remains a gold standard for assessing the integrity of barrier models, the typical handheld “chopstick” electrode TEER device requires removal of the cultures from the incubator to test each well individually, which can disrupt the cell layers. Additionally, variations in electrode positioning hinder reproducibility of TEER measurement due to the non-uniform electric field created by the chopstick electrodes (Benson et al., 2013; Giacomelli et al., 2017). Systems that integrate the electrodes directly into custom culture platforms (“EndOhm Chamber,” n.d.) or into microfluidic devices for microphysiological barrier modelling (Jeong et al., 2018; Maoz et al., 2017; van der Helm et al., 2016; Wang et al., 2017) address the challenges of manual handling and electric field uniformity to facilitate non-invasive, real-time monitoring. However, because TEER electrodes are placed on surfaces above and below the culture substrate, they measure bulk impedance of everything between the electrodes, not just the cell monolayer. As such, TEER measurement of cell monolayer barrier function is confounded in culture systems that contain multiple cell types or biomaterials (e.g., hydrogels), including most BBB models and microphysiological models.

An alternative to TEER is electrical cell-substrate impedance sensing (ECIS). ECIS applies the same principle as TEER except the electrodes are integrated into the substrate on which the cells are grown. This integration into the substrate allows cells to attach and proliferate directly on the electrodes, enabling more localised and sensitive impedance measurements specifically of the cell barrier compared to TEER. Furthermore, depending on the electrode placement and size, ECIS devices enable impedance measurements at higher electrical frequencies (e.g., 40 kHz), which can be used to determine additional characteristics of the cell layer (i.e., cell proliferation, cell viability, cell function, etc.) (Xu et al., 2016). The major disadvantage with current ECIS systems, however, is the limited compatibility of cell culture substrates with ECIS electrodes; current ECIS devices resemble standard printed circuit boards and require electrodes to be printed on either glass or stiff plastic. In order to implement ECIS capabilities in porous membrane systems typical for barrier modeling, alternative electrode printing strategies are required; this has proven to be challenging, forcing the use of expensive and custom-made devices (i.e., using e-beam evaporation to deposit gold onto the membrane (Gangopadhyay et al., 2017; Ramiah Rajasekaran et al., 2020)) that are impractical for large scale commercial manufacture. Moreover, these alternative printing strategies can result in high electrode impedance values that can drown out ECIS measurements.

To address this unmet need, we developed a cost-effective method to adapt ECIS technology to porous substrate-based *in vitro* models. Our approach enables high fidelity patterning of electrodes on porous membranes that can be incorporated into well plates of various sizes. We demonstrate that porous membrane ECIS (PM-ECIS) provides excellent cell biocompatibility with mono- and co-culture set ups, and sensitive, real-time measurement of changes in endothelial cell barrier impedance with cell growth and barrier disruption. Finally, we validate the system by direct correlation of PM-ECIS impedance and permeability coefficients obtained from molecular tracer permeability assays. Integration of ECIS into conventional porous cell culture inserts provides a versatile, sensitive, and automated alternative to TEER to measure barrier function in vitro.

## 2. Materials and Methods

### 2.1 Membrane electrode and device fabrication

Porous culture membranes were patterned with electrodes by heat-bonding gold leaf to the membranes (Figure 1A). Briefly, a stack consisting of (bottom to top): a flat glass sheet; polyethylene terephthalate (PET) porous track etched membrane (Sterlitech); a 200 nm thick gold leaf (on transfer paper; Gold Leaf Supplies); and a non-stick Teflon sheet (8569K34, McMaster-Carr), were bonded in a hot embosser (AutoSeries-3889, Carver) for 10 minutes at 2.3 MPa and 180°C. After letting the stack cool to room temperature under pressure, the gold leaf transfer paper was removed, and the stack placed back into the hot embosser to continue the bonding process for another 10 minutes at 9.2 MPa and 200°C. Finally, the stack was left under pressure to cool to room temperature. The initial thermal bonding step allowed for easy removal of gold leaf transfer paper, whereas the second bonding step fully bonded the membrane to the gold, leaving a smooth gold surface. A sheet of dry film photoresist (MonkeyJack) was then bonded to the gold surface using a laminator (ASC365, Thermal Laminating) at room temperature. Using a photomask, the negative pattern was exposed to UV for 10 seconds (OmniCure S2000, Lumen Dynamics) and developed in 1% Na_2_CO_3_ solution. The gold was etched using Gold Etchant TFA (Transene Company Inc. Danvers, MA) for 8 minutes (visually inspected to ensure all unwanted gold was removed) and washed with distilled water. The substrate was briefly washed with acetone to remove only the residual TFA etchant and then rinsed with distilled water. The remaining photoresist was removed using 0.5 M NaOH, after which the substrate was washed and left to soak in distilled water for at least 2 hours to ensure delamination of the membrane off the glass.

**Figure 1.**
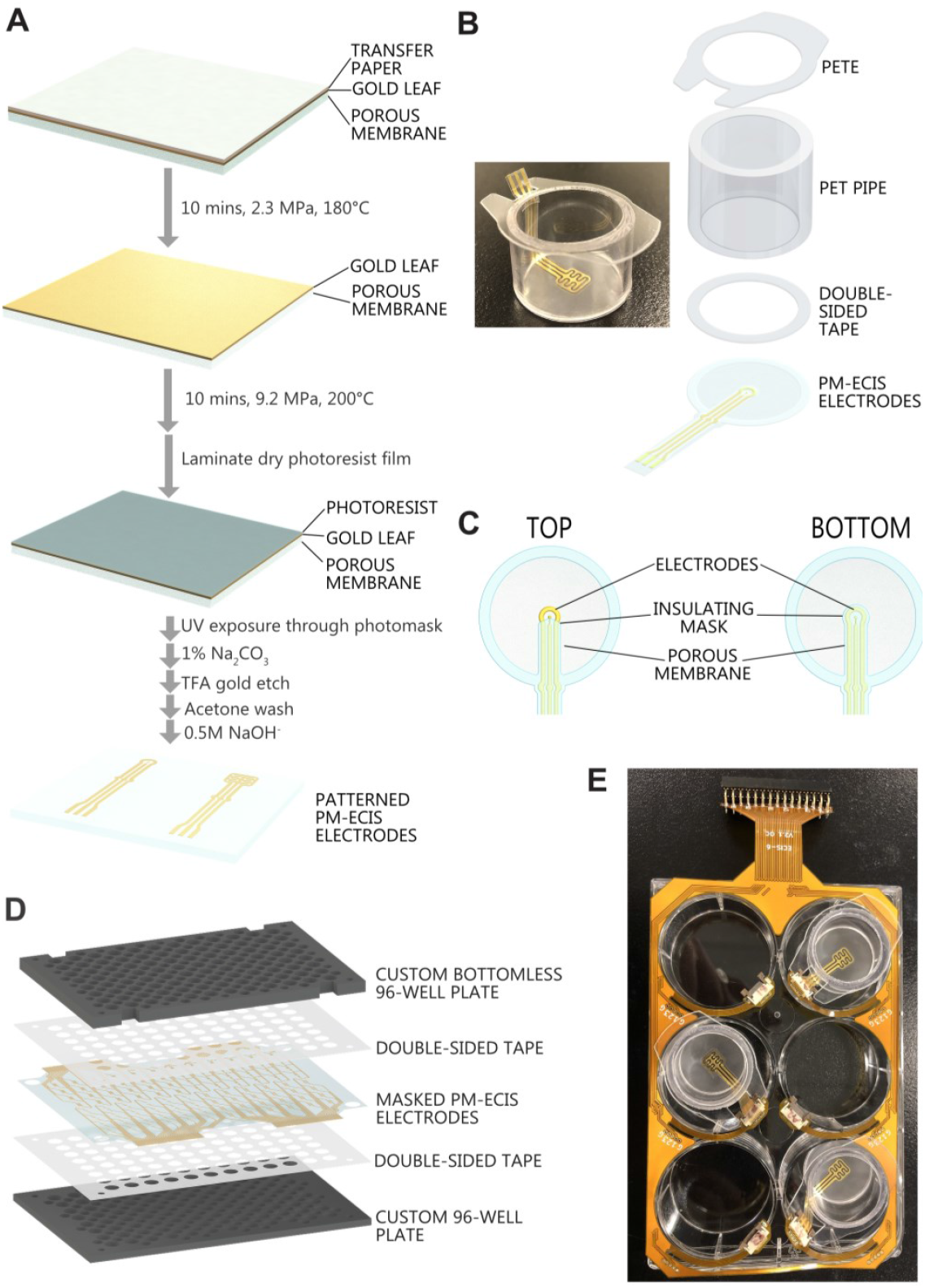
Porous membrane electrode fabrication and assembly into porous membrane ECIS (PM-ECIS) devices. (A) Gold leaf sheets were bonded to polyethylene terephthalate porous membranes using a pressured thermal bonding process, followed by photolithographic patterning and etching to form electrodes. (B) Fabrication of a custom 6-well “Transwell-like” insert incorporating PM-ECIS. (C) Application of electrical insulating masks on both sides of the porous membrane electrodes. (D) Fabrication of a custom 96-well device incorporating PM-ECIS. (E) Photograph of 6-well plate with three 6-well PM-ECIS inserts with flat flex cable gold finger connectors.

For cell culture, patterned PM-ECIS electrodes were cut using a laser cutter (VLS 3.5, Universal Laser Systems) for integration into custom-made 6-well inserts and 96-well devices. Inserts for 6-well plates were fabricated using PETG pipe (McMaster-Carr, #9245K37) with outer diameter = 1”, inner diameter = 7/8” cut to a height of 17.5 mm using a pipe cutter. A hot embossing step (140 kPa, 70°C. 3 min.) was used to create a smooth surface at the cut sites to facilitate subsequent assembly steps. Top frames for the inserts were created from laser cut polyester sheet (McMaster-Carr, #8567K92), and solvent bonded to the PETG pipe segments using SCIGRIP 4 Acrylic Cement (McMaster-Carr, #7517A2). A second hot embossing step (140 kPa, 70°C. 3 min.) further enhanced bonding between the polyester sheet and PETG pipe components. Finally, double-sided tape (ARSeal 90880, Adhesives Research) was used to secure PM-ECIS electrodes to the bottom face of the inserts; Fig. 1B summarizes assembly of the 6-well inserts.

Electrical insulating masks were laser cut from clear tape (Grand & Toy, #99842), aligned, and bonded on both sides of the membrane through cold lamination followed by hot embossing (2.15 kPa, 60°C, 2 min) to cover gold traces and restrict the sizes of electrodes (Fig. 1C).

PM-ECIS electrodes for 96-well devices were fabricated similarly to those in the insert embodiment. The electrodes were assembled with layers of laser cut double-sided tape and custom micro-milled polycarbonate 96-well plates (Fig. 1D).

Flat flex cable (FFC) gold finger connectors were added to the PM-ECIS electrodes to interface with a flex printed circuit board that connected the devices to the ECIS measurement system via an Arduino-operated multiplexer, allowing for measurements of up to 24 individual electrodes in parallel (Fig 1E); further details on the setup are provided in section 2.4.

Three electrode configurations were tested. For the 6-well plate inserts, either a single 500 μm diameter electrode (Fig. 2A,B; surface area 0.19635 mm2) or a six-finger electrode (Fig 2C,D; total surface area of 6 × 0.340175 mm2 = 2.04 mm2) was used. For the for 96-well plate, eight interdigitated electrodes were patterned (Fig. 2E,F; total electrode surface area of 4.933 mm2)

**Figure 2.**
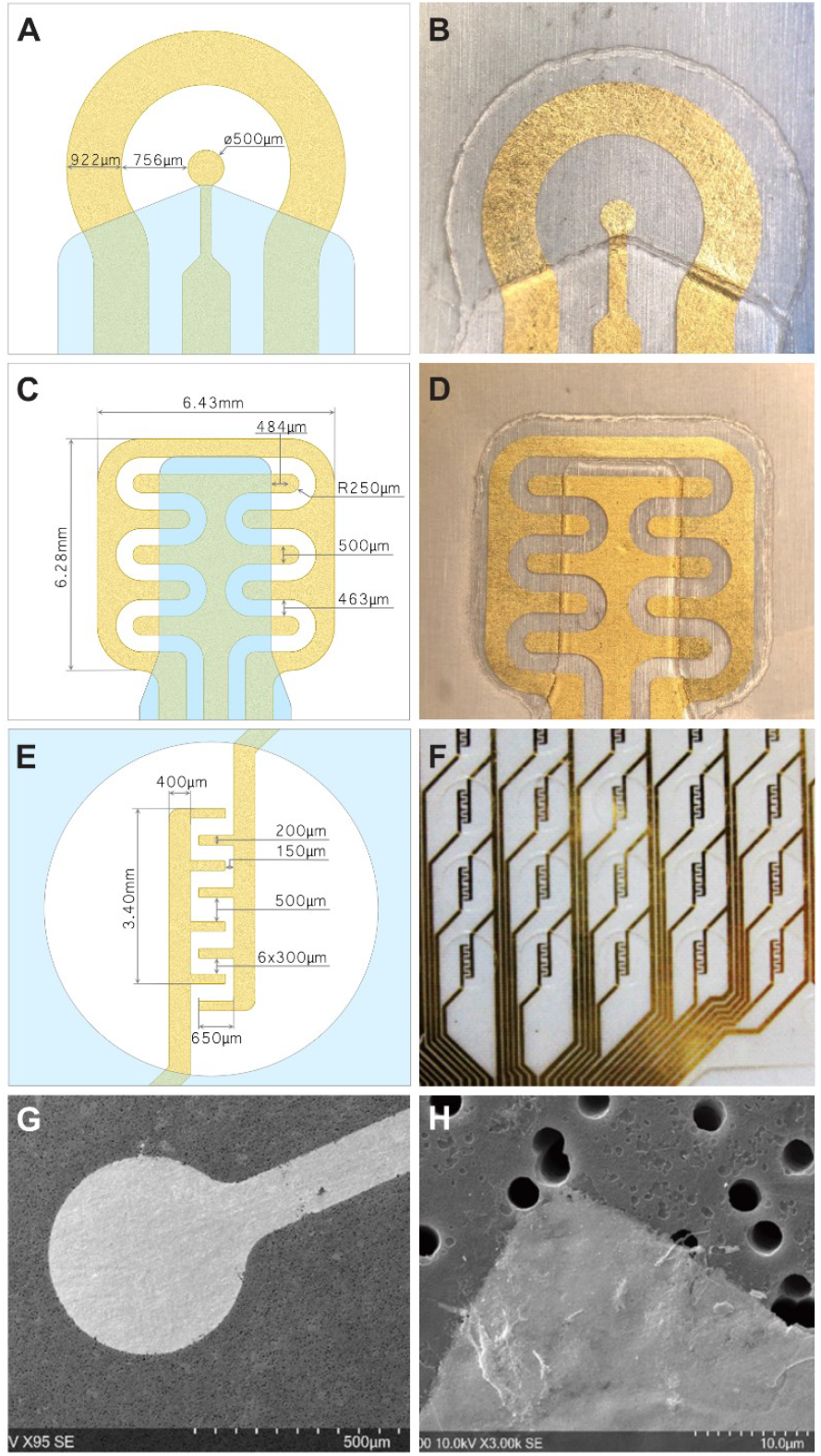
High fidelity patterning of different electrode designs on PM-ECIS devices. Schematics (left) and photographs (right) of (A, B) single electrode design for 6-well insert; (C, D) six-finger electrode design for 6-well insert; and (E, F) interdigitating eight-finger electrode design for 96-well plate. (G) Scanning electron micrograph of gold electrode patterned as the single electrode design on a porous membrane, showing high fidelity patterning and adhesion. (F) Scanning electron micrograph of interface of a patterned electrode and bare porous membrane, showing no deformation or clogging of the nominal 3 μm pores on the bare membrane.

### 2.2 Cell culture

Cells were cultured at 37°C and 5% CO_2_ in cell-specific media: primary human umbilical vascular endothelial cells (HUVECs, Lonza; up to passage 6) in EGM-2 (Lonza) culture medium kit; primary human brain microvascular endothelial cells (HBMVECs, Cell Systems -ACBRI 376) in complete classic medium (4Z0-500, Cell Systems) supplemented with CultureBoostTM (Cell Systems); immortalised HBMVECs (Boczula et al., 2021) in EGM-2MV microvascular endothelial cell growth medium (Lonza) supplemented with hygromycin-B (20 μg/mL); hCMEC/D3 BBB endothelial line (CELLutions Biosystems, Inc CLU512) cultured according to supplier’s directions in supplemented EBM-2 endothelial basal medium (Lonza); and normal human astrocytes (NHAs, Lonza) in Dulbecco’s minimum essential medium (DMEM), supplemented with N2 supplement (1%) and fetal bovine serum (10%). Media was changed every 48 hours in cell culture flasks, and every 24 to 48 hours in PM-ECIS devices.

To prepare PM-ECIS devices for cell culture, devices were plasma treated (PDC-001-HP, Harrick Plasma) for 3 minutes and UV sterilised for 2 hours (on both sides). Fibronectin (100 ug/mL) was then added to the apical side (with electrodes), and Geltrex (100 ug/mL) to the basolateral side for devices used in co-culture experiments, for 30 mins at RT to promote subsequent cell adhesion. HUVECs were seeded at 100,000 cells/cm2 or 250,000 cells/cm2 as indicated; primary HBMVECs were seeded at 5,000 and 30,000 cells/cm2 for permeability assays; and immortalized HBMVECs were seeded at 50,000 cells/cm2. For co-culture, astrocytes were seeded on the basolateral side of the ECIS device membrane at 50,000 cells/cm2 one day prior to endothelial cell seeding.

### 2.3 SEM imaging

Cell-free devices were imaged by scanning electron microscopy (Hitachi S-2500). To prepare for imaging, devices with cells were fixed with 4% paraformaldehyde for 2 hours and gradually dehydrated in 30%, 50%, 70%, 95% and 100% ethanol with 30 min incubation period for each step. The substrates were then sputtered with platinum (Polaron SC515 SEM coating system) and imaged.

### 2.4 Cell staining

For immunostaining, cells were fixed in methanol for 10 min at -20°C, washed three times with PBS, and blocked with 3% BSA (Sigma-Aldrich, 10735086001) for 20 min at 37°C. Cells were stained with primary antibodies (1:100) diluted in 3% bovine serum albumin in PBS for 1 hour at 37°C. Following incubation, cells were washed three times with PBS and blocked using 10% goat serum (Sigma-Aldrich, G9023) in PBS for 1 hour at room temperature. Secondary antibodies (1:200) diluted in 10% goat serum were applied for 1 hour at room temperature. After washing three times with PBS, nuclei were stained with 1:1000 Hoechst 33342 (Sigma-Aldrich, B2261) for 5 min at room temperature. Primary antibodies were mouse monoclonal ZO-1 (Thermo Fisher Scientific, 33-9100) and rabbit polyclonal GFAP (ab7260; Abcam); goat anti-mouse IgG (H+L) Alexa Fluor 488 (A32723, Thermo Fisher Scientific) and goat-anti rabbit IgG (H+L) Alexa Fluor 568 (A11036, Thermo Fisher Scientific). For viability staining, cells were stained using a Live/Dead kit (Invitrogen, USA, cat# L3224) according to the supplier’s protocol. Image acquisition was performed using an Olympus FV3000 confocal laser scanning microscope or Olympus BX51 inverted fluorescence microscope as indicated.

### 2.5 ECIS data collection

A lock-in amplifier (LIA) (SR850, Stanford Research Systems) was used to simultaneously generate AC current and read impedance data from the devices. An Arduino-operated multiplexer was used to switch between devices. Using a 1 V rms input voltage, in series with a 1 MΩ resistor, electrodes (typ. 500 μm dia.) were stimulated at frequencies of 400, 4000 and 40000 Hz, and any voltage drops (at the locked frequencies) across the electrodes were sampled with the LIA every 15 or 60 mins. Prior to use, the devices without cells were left in media for at least 12 hours to record a stable cell-free impedance baseline for each electrode at each frequency.

### 2.6 Impedance data analysis

Parasitic impedance offset at t = 0 was removed from all measurements, and then normalised to their effective exposed electrode areas (measured optically using a SZ61 Olympus stereoscope) using the following inverse sum equation:

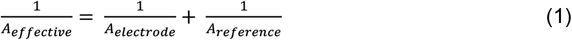

where A_electrode_ and the A_reference_ are the areas of the two electrodes used for impedance measurement.

Cell barrier resistance (R_b_), capacitance (C_m_), and basal barrier beneath the cells (α) were modeled and calculated using ECIS Core software (Applied Biophysics, v1.2.215.0 PC) using the raw impedance values taken at 400, 4000, and 40000 Hz.

### 2.7 Thrombin stimulation

For experiments in which the endothelial barrier was disrupted with thrombin treatment, cell culture media was replaced with fresh media containing 1 U/mL of thrombin from bovine plasma (Sigma) at the stimulus onset. Fresh media without thrombin was used as a vehicle only control.

### 2.8 Permeability assay and analysis

To measure barrier permeability, 1 mL of 50 μg/mL 10 kDa FITC-dextran in phosphate-buffered saline with calcium and magnesium (PBS+/+) was added to the apical side of a 6-well membrane PM-ECIS device, and 2 mL of PBS +/+ to the basolateral side. Volumes were chosen to match volume height and not induce pressure driven flow across the membrane. 100 μL samples were taken from the basolateral side (after pipette mixing) at 0, 5, 10, 20, and 30 minutes. 100 μL of PBS +/+ was replenished to each well after every sample. Permeability constants were calculated and modeled via the methods described by Hubatsch et al. (Hubatsch et al., 2007).

## 3. Results and Discussion

### 3.1 Gold leaf heat-bonding is a cost- and time-efficient alternative to metal deposition for membrane electrode fabrication

Fabrication methods such as metal evaporation (thermal and e-beam) and sputtering are the current industry-standard methods for depositing gold electrodes onto non-conductive surfaces due to their versatility in a wide range of applications (Henry et al., 2017; Sokolov et al., 2009). They are limited, however, by high material and time costs that make post-prototyping manufacture challenging. Here we developed a gold leaf transfer protocol (Fig. 1A) as a simple, cost-efficient alternative to metal deposition for manufacturing membrane electrodes for PM-ECIS. The bonding process relies on the membrane reaching its glass transition temperature and mechanically bonding to the gold leaf. By controlling the bonding temperature and pressure, bonding can be achieved without deformation or clogging of the pores in the PET membranes or compromising their integrity (Fig. 2G,H).

Heat-bonding pre-manufactured gold sheets directly to porous membranes has numerous advantages over traditional metal deposition protocols. In particular, heat-bonding gold leaf provides a flat continuous substrate that permits standard large scale photolithography techniques and results in improved conductivity over deposition methods. For example, we measured electrode resistivity of commercial roll-to-roll sputter-coated devices to be 3.8 × 10-8 Ω·m, whereas the resistivity of the same electrode pattern fabricated by gold leaf heat-bonding was 2.6 × 10-8 Ω·m; lower electrode resistance values are more desirable as they allow for higher sensitivity measurements. The leaf heat-bonding method also allows for the use of biocompatible gold alloys (silver and copper) that further improve electrical conductivity and electrode integrity. Finally, there is minimal material wastage with leaf heat-bonding, as the only gold lost during fabrication is the negative space between the electrodes. In evaporating or sputtering methods, the majority of the gold is wasted due to the low solid angle ratio between the exposed substrate and the rest of the machine. We estimate that for the PM-ECIS devices reported here, fabrication by standard thermal evaporation and photolithography techniques would be ∼4-fold more expensive and take ∼35% longer than gold leaf bonding.

### 3.2 PM-ECIS devices support and monitor endothelial cell adhesion, growth, and monolayer barrier formation

To test their biocompatibility, PM-ECIS devices were seeded with HUVECs. In 96-well PM-ECIS devices, HUVECs grew on the fibronectin-coated gold electrodes, tape masks, and PET membranes at similar densities and with similar viabilities and morphologies to HUVECs on standard tissue culture-treated polystyrene well plates (Fig. 3A-C). SEM imaging further confirmed HUVEC attachment to electrodes of different shapes (Fig. 3D,E), with cells spanning the interface between the electrodes and the bare membrane (Fig. 3D-F). Moreover, SEM imaging confirmed the electrodes stayed fully adhered to membranes after at least 4 days in culture (Fig. 3D-F).

**Figure 3.**
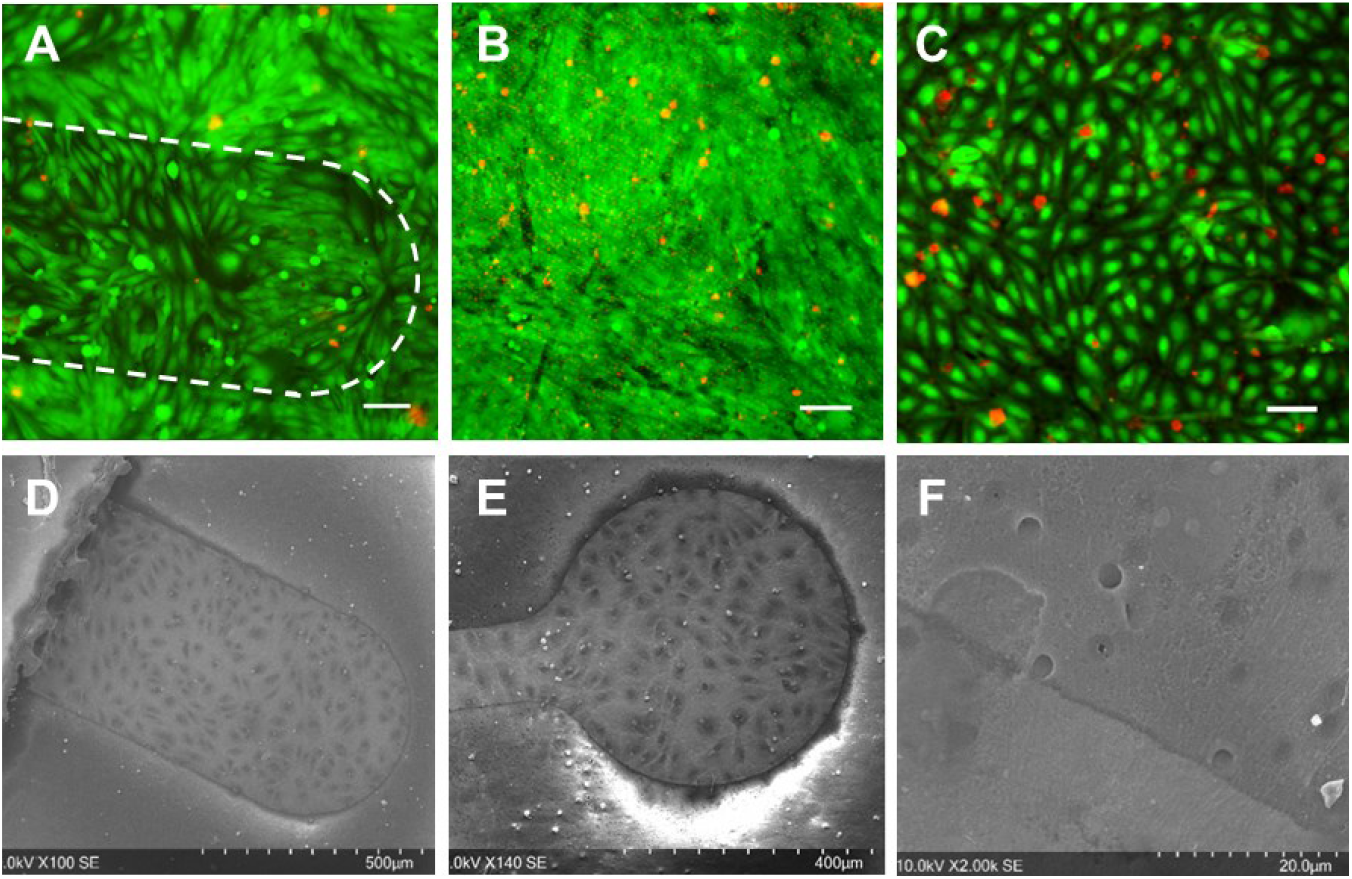
PM-ECIS devices support endothelial cell culture. Human umbilical vein endothelial cells (HUVECs) stained with calcein AM and ethidium homodimer to indicate live (green) and dead (red) cells, respectively, showed high viability when grown on (A) gold electrodes (indicated by white dotted line) and the adjacent porous membrane and (B) areas of the membrane with the tape mask, comparable to HUVECs grown on (C) standard tissue culture-treated polystyrene plates. Scanning electron micrographs of 4 day cultures confirmed HUVEC adhesion to and coverage of (D) finger-like and (E) round electrodes, with (F) the cell layer spanning the interface of the electrode and bare membrane, with no visible difference in cell morphology. Scale bars in A-C = 100 μm.

Electrical impedance was measured at 400 Hz, 4 kHz, and 40 kHz in 96-well devices over 4 days of culture, with cells initially seeded at 100,000 or 250,000 cells/cm2. Resistance at 4 kHz increased over the culture duration, indicating increasing cell coverage of the electrodes (putatively due to spreading and proliferation) and increasing barrier function (Fig. 4A). As expected, cells seeded at 250,000 cells/cm^2^ had greater resistance initially than cells seeded at 100,000 cells/cm^2^ (Fig. 4A). Resistance in the higher density cultures also plateaued by day ∼2.5, suggesting confluence and barrier formation, whereas resistance of the lower density cultures continued to increase throughout the four days (Fig. 4A). Cell-free controls showed no change in resistance over the culture period. Similar to traditional solid substrate ECIS systems (Applied Biophysics, n.d.; Gamal et al., 2018), events in cell culture, including cell micromotion (high frequency oscillations in resistance) and media changes (abrupt changes in resistance), were also observed in the electrode signal (Fig. 4A).

**Figure 4.**
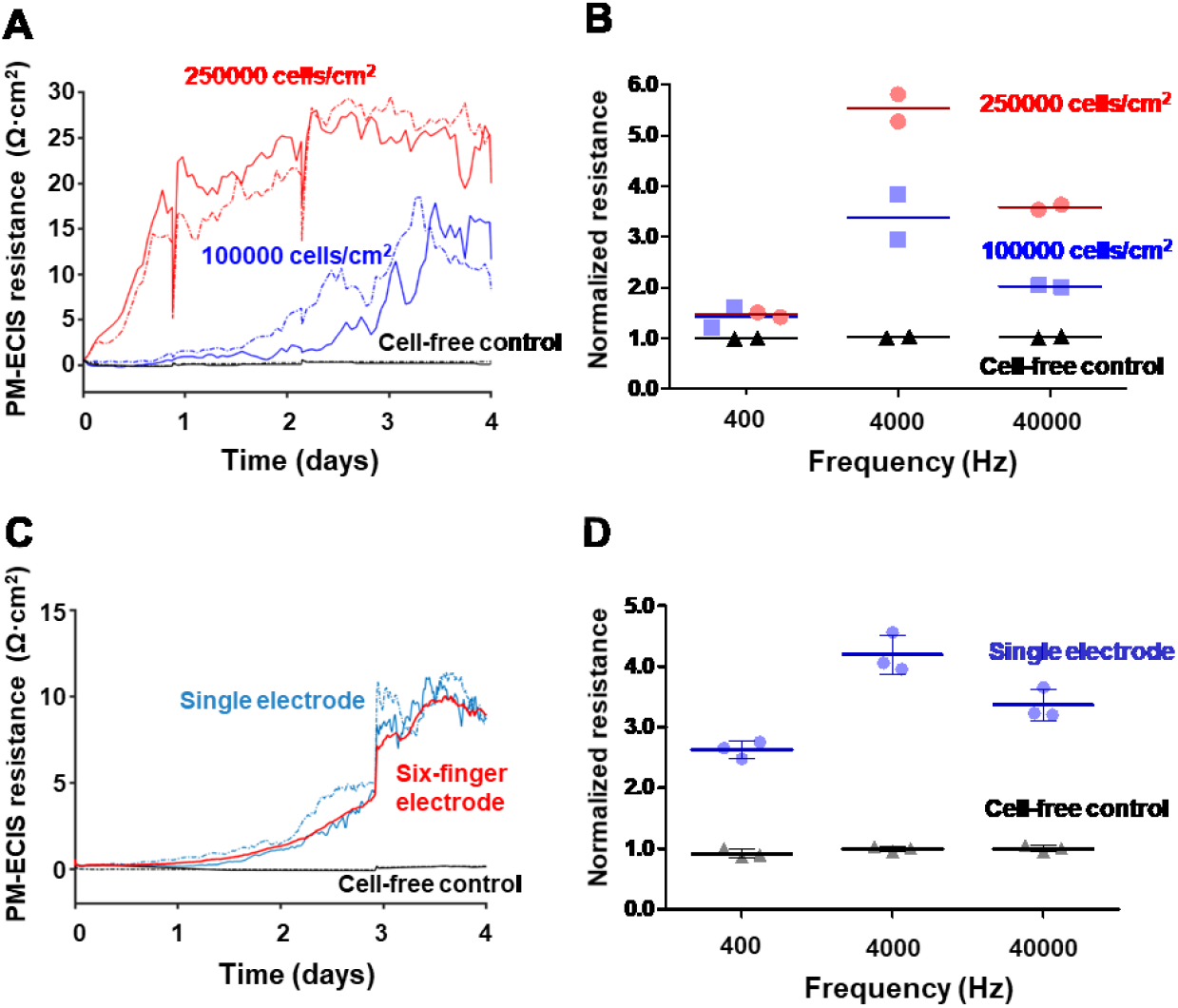
PM-ECIS measurement of endothelial monolayer resistance is sensitive to seeding density. (A) Representative traces of 4 kHz PM-ECIS measurement of human umbilical vein perivascular cells (HUVECs) initially seeded at densitites of 250,000 cells/cm^2^ (red lines) or 100,000 cells/cm^2^ (blue lines) in 96-well PM-ECIS devices. Dotted lines show traces from a separate device under the same conditions. Black lines are cell-free control devices. (B) PM-ECIS resistances normalized to cell-free controls across the three measured frequencies (400, 4000, 40000 Hz) at end of day 4 for the high (red) and low (blue) density cultures. (C) Representative traces of 4 kHz PM-ECIS resistances measured for HUVECs cultured in 6-well devices with either a single electrode (blue) or 6-finger electrode (red). Dotted lines show traces from a separate device under the same conditions. Black lines are cell-free control devices. (D) PM-ECIS resistances normalized to cell-free controls across the three measured frequencies (400, 4000, 40000 Hz) at end of day 4 for the 6-well single electrode devices with (blue) and without (black) HUVECs. N = 3 devices.

The final cell monolayer resistances at 4 kHz were 10-30 Ω·cm^2^ (Fig. 4A,B), consistent with values from commercial solid substrate ECIS systems (Keese et al., 2002; Stolwijk et al., 2015). Also similar to trends observed with commercial solid substrate ECIS systems (Giaever and Keese, 1991; Stolwijk et al., 2015; Szulcek et al., 2014), cell monolayer resistance was greater at 4 kHz than 400 Hz or 40 kHz and was greater in the higher density cultures (Fig. 4B). HUVECs grown on the 6-well PM-ECIS inserts displayed a similar behavior to cells grown in 96-well plate devices. The resistance of cells seeded at 100,000 cells/cm2 gradually increased over time in culture to ∼10 Ω·cm^2^ by day 4, whereas cell-free controls showed no change (Figure 4C). Furthermore, similar trends were observed for the two electrode designs tested in these experiments, although the six-finger electrode with larger area produced a steadier ECIS signal than the single 500 μm diameter electrode (Fig. 4C); this was similar to commercial ECIS devices with comparable electrode designs (Applied Biophysics, n.d.; Stolwijk et al., 2015). Cell monolayer resistance peaked at 4 kHz for both electrode designs (Fig. 4D).

### 3.3 PM-ECIS devices support and monitor cell co-culture

A significant advantage of PM-ECIS devices over traditional solid substrate ECIS devices is that porous membranes enable co-culture, as is typical for BBB modeling. To test our 6-well PM-ECIS inserts for this application, immortalized BBB endothelial cells and primary human astrocytes were cultured separately or together on the devices. Both cell types grew well when cultured separately on the devices, with HBMVECs forming confluent layers with junctional expression of ZO-1 (Fig. 5A) and astrocytes spreading on the membranes and expressing GFAP (Fig. 5B). Similarly, when co-cultured with the HBMVECs on the apical surface of the membrane (where the electrodes were patterned) and the astrocytes on the basal surface, both cell types grew as monolayers on the membranes and expressed their phenotypic markers (Fig. 5C).

**Figure 5.**
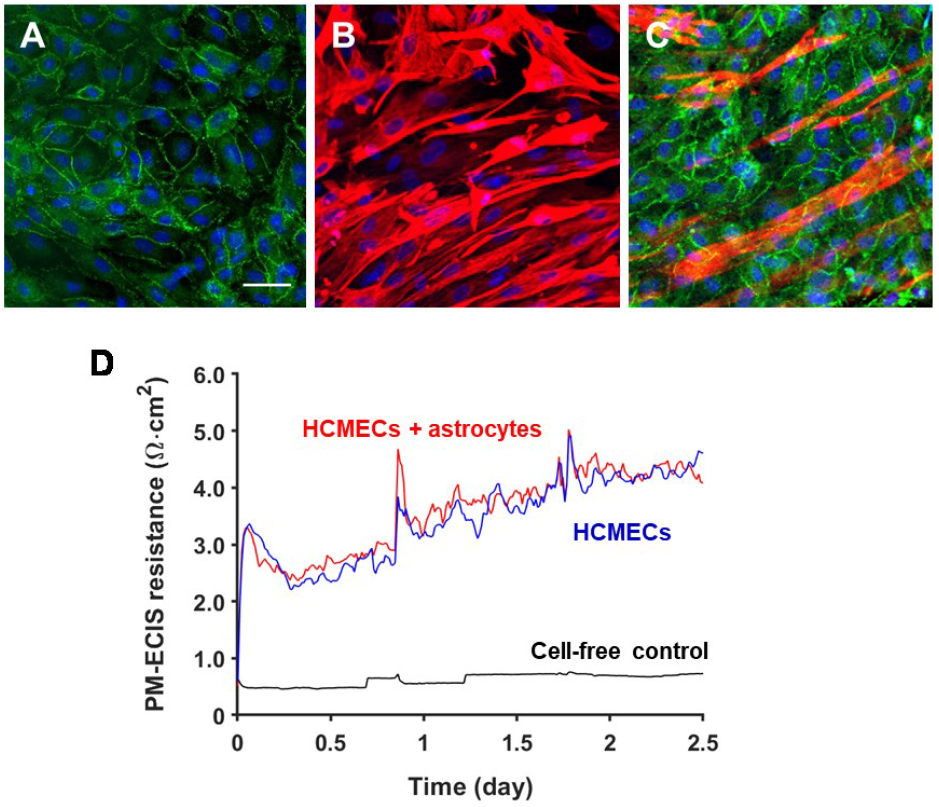
PM-ECIS devices support co-culture of endothelial cells and astrocytes. (A) Immortalised HBMVECs grown in monoculture on the apical side of 6-well PM-ECIS devices after 4 days in culture, stained for ZO-1 (green) and nuclei (Hoescht dye; blue). Scale bar = 100 μm. (B) Human astrocytes grown in monoculture on the underside of the PM-ECIS device, stained for GFAP (red) and nuclei (blue). C) Confocal maximum intensity projection of apical side HBMVECs co-cultured with basolateral side astrocytes, demonstrating phenotypic ZO-1 (green) and GFAP (red) staining, respectively. Nuclei stained by Hoescht in blue. (D) PM-ECIS resistance measurement at 4000 Hz for 2.5 days of culture, showing endothelial layer resistance was unaffected by the presence of co-culture astrocytes on the opposite of the membrane.

ECIS resistance was measured for the monoculture and co-culture conditions in the 6-well, single electrode devices. As expected, brain endothelial cells (HCMEC/D3 line in Fig. 5D) grown on the apical surface of the membrane alone or in co-culture with astrocytes on the basal surface showed increasing 4 kHz PM-ECIS resistance over 2.5 days in culture, with typical minor fluctuations reflecting local cell movement on the single electrode (Fig. 5D). Importantly, the resistance in co-cultures with astrocytes on the basal surface of the membrane was similar to that with monoculture, reflecting that the cells on the basal membrane surface do not contact the electrodes. The ability of membrane ECIS to isolate measurement of barrier function to only the barrier-forming cell monolayer is an important feature that is not possible with TEER.

The lack of an effect of co-culture of primary astrocytes on PM-ECIS resistance of the immortalized HCMEC/D3 line used here was consistent with previous studies by Garcia-Salvador *et al*. (2020) noted that neither primary rat astrocytes nor their conditioned medium improved hCMEC/D3 barrier integrity, whereas *immortalized* rat astrocytes and their conditioned medium did; the authors suggested this reflected the immortalized astrocyte line maintaining its phenotype more effectively than primary astrocytes in vitro. Nonetheless, culture of some brain endothelial cell lines with either astrocyte-conditioned media (Rubin et al., 1991; Siddharthan et al., 2007) or directly with astrocytes (Boveri et al., 2005; Cecchelli et al., 1999; Perrière et al., 2007; Takeshita et al., 2014) can increase TEER barrier resistance, although this effect is not seen with all cell lines (Eigenmann et al., 2013; García-Salvador et al., 2020). We would expect robust paracrine signaling between the apical and basal compartments of the PM-ECIS devices, as the membrane porosity is retained over the bulk of the membrane. Direct cell contact via cell protrusions through the membrane pores would also be possible (Alvarez et al., 2013), except in regions with electrodes. Blockage of direct cell-cell contact could be a limitation of the PM-ECIS devices for certain applications; further investigation would be needed to determine if the minimal effect of astrocyte co-culture on PM-ECIS resistance observed in this study was due to minimal contact locally or to other factors, including cell line, time in culture, and cell densities.

### 3.4 PM-ECIS devices enable measurement of pericellular resistances and cell membrane capacitance

The total impedance between ECIS electrodes can be modeled as network of resistive and capacitive loads resulting from both the cell layer and non-cell impedances (i.e., capacitive interactions between electrodes and media) (Giaever and Keese, 1991). Non-cell related parasitic impedances can be isolated and removed by measuring a baseline for each electrode pair before adding cells, as well as having a parallel cell-free control to measure electrode impedance drift over time. Pericellular resistance (R_b_) across the tight junctions, a measure of true barrier integrity, can then be approximated via the impedance transfer function developed by Giaver and Keese (Giaever and Keese, 1991) by recording resistive and capacitive impedance components across a range of frequencies. This resistance is a linear measurement of the free space between the cells where small molecules can simply diffuse. The model also determines values for cell membrane capacitance (C_m_) and basal pericellular impedance (α) between the cell layer and the electrodes; α is a function of the ratio between the average cell radius and the distance between the cell layer and the electrodes.

R_b_, α, and C_m_ values were measured in HUVEC monolayers on the 6-well PM-ECIS inserts (Figure 6). As expected, R_b_ rapidly increased with cell monolayer formation to a peak value of ∼6 Ω·cm^2^ 60 hours post-seeding (Fig. 6A). Basal pericellular impedance, α, peaked at ∼8 Ω^1/2^·cm during the first day of culture, and then gradually decreased to ∼6 Ω^1/2^·cm after 60 hours (Fig. 6B), reflecting the reduction in the cells’ sizes as they flattened and packed together to form a confluent monolayer. The values measured here for R_b_ and α were similar to those reported for solid substrate ECIS devices: 3.0 – 3.9 Ω·cm^2^ and 5.3 -5.4 Ω^1/2^·cm for R_b_ and α, respectively (Pannekoek et al., 2013; Robilliard et al., 2018; Stolwijk et al., 2015; Szulcek et al., 2014). We observed initial values of C_m_ of ∼1.4 μF/cm^2^, which is similar to that observed with commercial solid substrate ECIS systems (1.1 – 1.4 μF/cm^2^) (Pannekoek et al., 2013; Robilliard et al., 2018; Stolwijk et al., 2015). C_m_ rapidly decreased over the first hours of culture coincident with the increases in R_b_ and α as the cells spread and became confluent, and then plateaued ∼0.3 μF/cm^2^ (Fig. 6C). These trends also match those reported using solid substrate ECIS with epithelial cells (Ebrahim et al., 2022) and brain endothelial cells (Kho et al., 2017; Robilliard et al., 2018).

**Figure 6.**
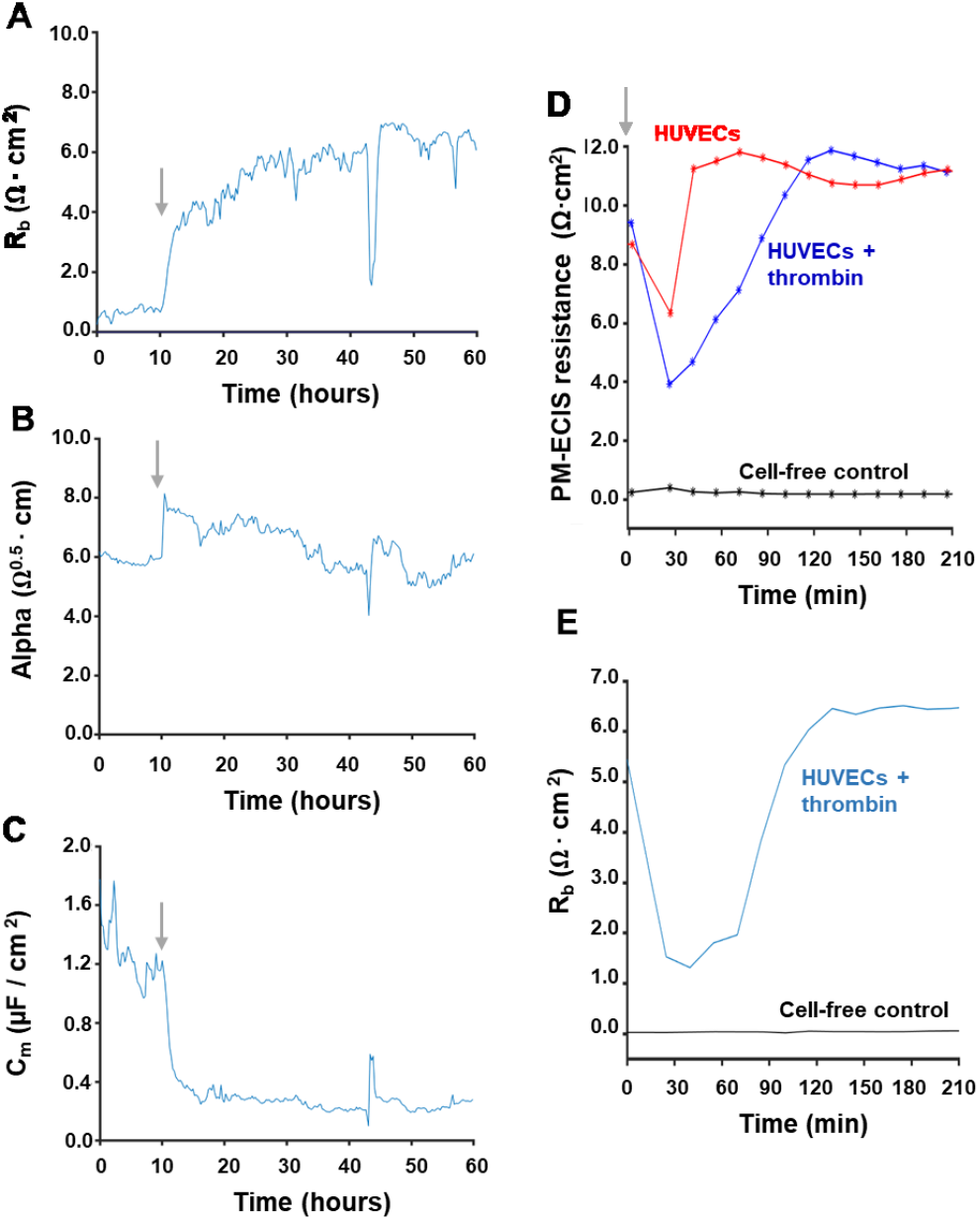
Pericellular resistances and cell membrane capacitance can be derived from PM-ECIS measurements. Human umbilical cord perivascular cells (HUVECs) grown in 6-well devices were treated with serum-containing medium hour 10 (grey arrows) and monitored continuously by PM-ECIS. The network model developed by Giaver and Keese (1991) was applied to derive (A) cell-cell pericellular resistance, R_b_; (B) cell-electrode pericellular resistance, α; and C) cell layer membrane capacitance, C_m_. (D) 4kHz PM-ECIS resistance measurements of HUVECs treated a t=0 (grey arrow) with media only (HUVECs) or media + thrombin (HUVECs + thrombin) to cause barrier distruption. (E) Calculated pericellular resistance, R_b_, for thrombin-disrupted HUVEC layer.

To confirm that the increase in measured resistance (and R_b_) was due to cell barrier formation, cells were treated with thrombin to disrupt cell-cell junctions. Indeed, the addition of thrombin resulted in an acute resistance drop (Figure 6D), indicating a temporary disruption in pericellular resistance, estimated by the Rb parameter (Figure 6E); similar trends were observed in commercial solid substrate ECIS systems (Andreeva et al., 2010; Szulcek et al., 2014). Control cultures in which only fresh media was added (vehicle control) also demonstrated a small drop in resistance (Fig. 6D); this is typical with media changes (e.g., Fig. 4A) and recovery was much quicker (∼15 min) than with thrombin treatment (∼120 min).

### 3.5 PM-ECIS devices allow for simultaneous investigation of permeability coefficients and cell layer resistance

An advantage of PM-ECIS is that the porous membrane allows for simultaneous measurement of permeability and electrical resistance. To demonstrate this and test their correlation, we seeded primary HBMVECs on PM-ECIS devices at densities of 5,000 cells/cm^2^ or 30,000 cells/cm^2^ and cultured them for 4.5 days. Barrier resistances at 400 Hz, 4 kHz, and 40 kHz were monitored by PM-ECIS throughout the culture period, and FITC-dextran permeability assays were performed at days 1, 2, 3, and 5. Differences in membrane coverage between the two seeding densities and with time in culture were reflected in higher PM-ECIS resistances for the higher density cultures at all frequencies and more rapid increases in resistances as the cells proliferated, spread, and formed barriers (Fig. 7A-C). The differences between the cultures initially seeded with 5,000 cells/cm^2^ or 30,000 cells/cm^2^ were also evident in the FITC-dextran permeability measurements: the day 4.5 permeability constants were 5.57 × 10^−3^ ± 0.912 × 10^−3^ cm/min and 1.54 × 10^−3^ ± 0.593 × 10^−3^ cm/min, respectively. Notably, we observed large drops in PM-ECIS resistances after each permeability measurement, followed by lengthy lags recovery periods, which were sometimes incomplete (Figs. 7A-C). This reflects the known disruptive effect of dextran on endothelial viability and function (Rouleau et al., 2010), and highlights the advantage of PM-ECIS for sensitive real-time measurement of barrier integrity without disruption of cultures. Nonetheless, we observed significant negative correlations between FITC-dextran permeability coefficients and PM-ECIS resistances at each of 400 Hz (Fig. 7D; r = -0.835, p=0.019), 4 kHz (Fig 7E; r = -0.836, p=0.019), and 40 kHz (Fig. 7F; r = -0.795, p=0.033). Strong correlation between PM-ECIS and a tracer permeability assay is notable as it suggests that barrier function is similar between cells grown on the electrode (as measured by PM-ECIS) and directly on the bare membrane (as measured by a tracer assay), consistent with our observations of uniform cell coverage (Fig. 3).

**Figure 7.**
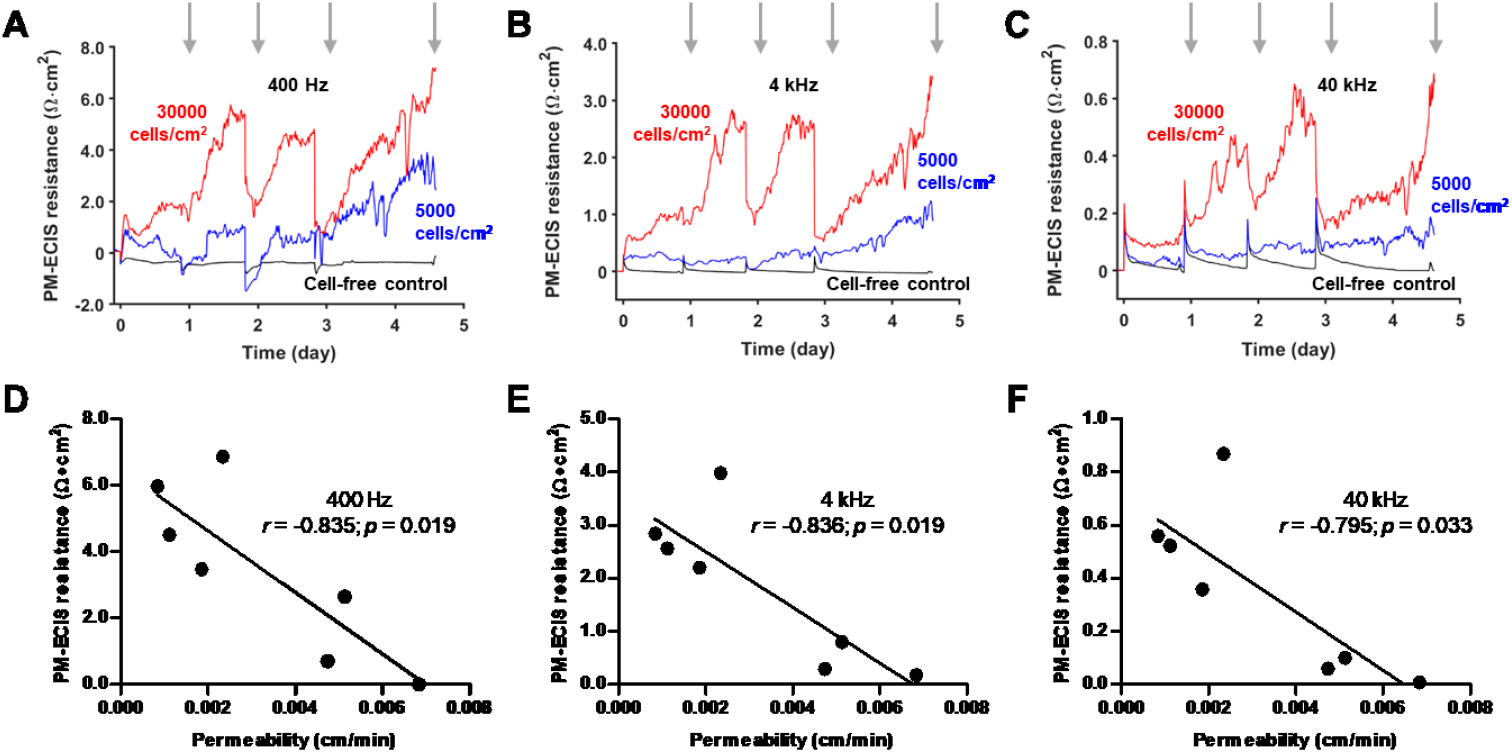
Endothelial monolayer resistances measured by PM-ECIS correlate with permeability measurements. Primary HBMECs were seeded on 6-well PM-ECIS devices at initial (day 0) seeding densities of 5,000 cells/cm^2^ or 30,000 cells/cm^2^, and 10 kDa FITC-dextran permeability and PM-ECIS measurements performed on day 1, 2, 3, and 4.5 post-seeding (indicated by grey arrows in panels A-C). PM-ECIS was measured continuously throughout the experiment, with measurements made at (A) 400 Hz, (B) 4 kHz, and (C) 40 kHz. Final (day 4.5) PM-ECIS resistances measured at (D) 400 Hz, (E) 4 kHz, and (F) 40 kHz were significantly correlated with FITC-dextran permeabilities measured in the same devices. N = 3 devices initially seeded at 5,000 cells/cm^2^ and N = 4 devices initially seeded at 30,000 cells/cm^2^.

## 4. Conclusions

ECIS is a powerful method for non-invasive, real-time monitoring of cell behavior and viability in vitro, but has largely been restricted to solid culture substrates. To enable ECIS measurements on standard porous membrane culture inserts, we developed a cost-effective method to deposit gold electrodes on porous membranes with high fidelity. We showed that PM-ECIS provides sensitive, real-time measurement of changes in cell barrier impedance with cell growth and barrier disruption in both monoculture and co-culture set ups, with direct correlation with permeabilities measured by a molecular tracer assay. In total, PM-ECIS is a versatile, high sensitivity alternative to current methods to measure barrier function in vitro, including TEER and molecular tracer assays.

## 5. Acknowledgements

We thank Applied Biophysics for providing advice and software for the network model analysis; Prof. Tara Moriarty and Anna Boczula (University of Toronto) for providing the immortalized HBMVECs; and Hayley Yap for assistance with figure creation. This work was funded by a CQDM Explore grant, the Natural Sciences and Engineering Research Council of Canada (CRDPJ 531083-18), and the Ontario Centre of Excellence (VIP II 29253).

## Author contributions

**Oleg Chebotarev:** Conceptualization; Formal analysis; Investigation; Methodology; Visualization; Writing - original draft; **Alisa Ugodnikov:** Data curation; Visualization; Writing - review & editing; **Craig Simmons:** Conceptualization; Formal analysis; Funding acquisition; Project administration; Supervision; Visualization; Writing - review & editing.

